# PlantGeneAnn: a high-accuracy *ab initio* plant gene annotation tool fine-tuned from a genome foundation model

**DOI:** 10.64898/2026.06.25.733695

**Authors:** Qizhe Zhang, Zhengyang Zhang, Kepeng Lin, Jing Wang, Kaixuan Deng, Wei Xu, Qing-Yong Yang, Xuehai Hu

## Abstract

Structural annotation remains a bottleneck for translating the growing catalog of high-quality plant genomes into biological insights, as current pipelines rely on RNA-seq and homologous protein evidence that are often unavailable for non-model species. Existing *ab initio* tools are constrained by model scale and perform poorly in complex plant genomic regions. We present PlantGeneAnn, a plant-optimized *ab initio* gene annotation tool fine-tuned from a genome foundation model. It combines a Mamba-MoE DNA encoder with a U-Net-style embedding decoder and a semi-Markov decoding pipeline. Trained on 40 angiosperm species, PlantGeneAnn establishes a new state-of-the-art (SOTA) across a stringent 20-species benchmark, significantly outperforming Helixer, Tiberius, and ANNEVO at exon-, intron-, and locus-levels, with mean F1 scores of 0.856, 0.880, and 0.707, respectively. It also enables efficient customized fine-tuning: LoRA fine-tuning increased the locus-level F1 of the out-of-distribution bryophyte *Physcomitrium patens* from 0.581 to 0.736, with performance plateauing at 40% of the fine-tuning data. Notably, PlantGeneAnn requires substantially lower peak memory usage than other baseline tools, allowing annotation of large plant genomes on standard personal workstations. PlantGeneAnn thus provides accurate, hardware-friendly ab initio annotation for diverse plant genomes, particularly newly assembled genomes, population-scale projects, and non-model plants.

## Introduction

Long-read sequencing, telomere-to-telomere assembly, and population-scale sequencing are rapidly expanding the catalog of high-quality plant genomes. However, genome sequences can support biological discovery only when genes, exons, and coding sequences (CDSs) are accurately defined. Such structural annotations provide the foundation for gene function studies, comparative genomics, and crop improvement (Michael, 2026).

Current gene annotation pipelines integrate RNA-seq with homologous protein evidence. Although they produce accurate gene models when external evidence is abundant, their performance depends on comprehensive sampling across tissues and developmental stages and on the evolutionary proximity of reference species (Ji et al., 2026). Consequently, genes with restricted expression, weak homology, or lineage-specific functions may be poorly annotated, particularly for newly assembled genomes, population-scale projects, and non-model plants.

When external evidence is limited or unavailable, *ab initio* gene annotation becomes essential. Over the past two decades, this field has progressed from hidden Markov model (HMM)-based tools, such as AUGUSTUS (Stanke and Waack, 2003), to deep-learning methods, including Helixer (Holst et al., 2026), Tiberius (Gabriel et al., 2026), and ANNEVO (Zhang et al., 2026), and more recently to genome foundation model (GFM)-based approaches, such as Nucleotide Transformer-v3 (Boshar et al., 2025). However, they all have some shortcomings: HMM-based tools require species-specific training or auxiliary evidence; earlier deep-learning methods, constrained by limited capacity, struggle with complex plant genomic regions enriched in repetitive sequences, transposable elements, and lineage-specific gene family expansions. GFM-based tools, a new AI-era solution, have not reached consensus on whether they can surpass previous strategies and become optimal for gene annotation, especially in plants. Here, we developed **PlantGeneAnn** (**Plant Gene Ann**otator), a plant-optimized *ab initio* annotation tool based on our previously published GFM, PlantBiMoE (Lin et al., 2025). Benchmarking against existing tools in prediction performance and computational cost shows PlantGeneAnn is more robust and accurate, achieving a state-of-the-art (SOTA) in structural annotation accuracy for plant genomes.

## Results

PlantGeneAnn consists of three major modules: genome slicing, sequence modeling, and gene structure decoding (Figure 1A). The query genome is first divided into 40k-bp fragments and then processed by a model comprising a DNA encoder, an embedding decoder, and finally reach the output heads (Supplemental Figure 1). The model was trained on 105 billion tokens from 40 angiosperm species (Figure 1B), with the DNA encoder initialized from PlantBiMoE (Figure 1A). The embedding decoder adopts a U-Net-style architecture with a Transformer bridge to model long-range dependencies (Supplemental Figure 1). The output heads generate probabilities for 15-state labels on each strand, which are further decoded by a biological knowledge-constrained semi-Markov model (SMM) to output gene structure (Supplemental Figure 2).

**Figure 1.**
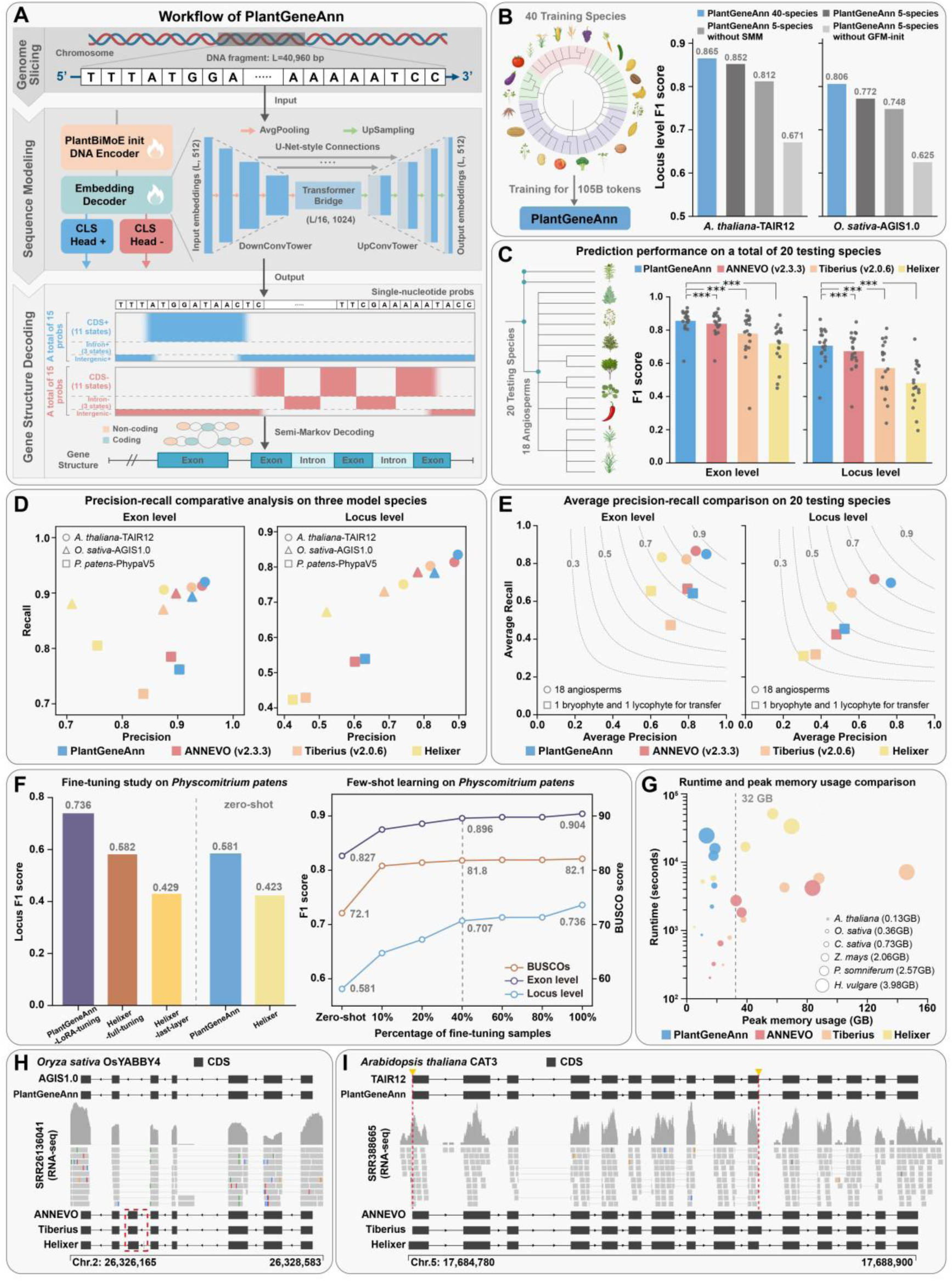
Workflow, *ab initio* annotation performance, customized fine-tuning analysis, computational resource comparison, and annotation examples of PlantGeneAnn. (**A**) Schematic of the PlantGeneAnn workflow, including genome slicing, sequence modeling, and gene structure decoding. **(B)** Ablation analysis of key model components. Locus-level F1 scores on Arabidopsis and rice are shown for the complete PlantGeneAnn model (trained on 40 species, 40-species), the ablation baseline (trained on five species, 5-species), and ablated models without SMM post-processing or without GFM-based weight initialization. All ablated variants were trained in the five-species setting. **(C)** Exon– and locus-level F1 scores for PlantGeneAnn and three baseline tools across 20 test species. Bars indicate means and dots indicate individual species. Asterisks indicate significant differences of PlantGeneAnn versus each baseline tool (two-sided paired Wilcoxon signed-rank tests; *** P < 0.001). **(D)** Precision–recall analysis in Arabidopsis, rice, and *Physcomitrium patens*. **(E)** Mean precision–recall analysis across 20 test species. Circles indicate 18 angiosperms, and squares indicate a bryophyte and a lycophyte used to assess cross-lineage transferability. **(F)** LoRA fine-tuning and few-shot learning analyses on *Physcomitrium patens*, showing F1 scores and BUSCO completeness under different fine-tuning strategies and sample sizes.**(G)** Comparison of annotation runtime and peak system memory usage across six species with different genome sizes. Bubble size indicates genome size. **(H)** Gene structure comparison at the rice *OsYABBY4* locus. PlantGeneAnn reproduced the reference structure, whereas all three baseline tools predicted an additional exon between exons 5 and 6. **(I)** Gene structure comparison at the Arabidopsis *CAT3* locus. PlantGeneAnn correctly recovered the complete gene structure, whereas other baseline tools misidentified either the start codon or the splice donor site.

To determine how each component within the model architecture contributes to performance, we performed ablation analyses on a smaller dataset comprising five species (Supplemental Table 1). Results from Arabidopsis and rice showed that multi-species training (40-species), SMM-based post-processing, and GFM weight initialization all improved performance (Figure 1B). In addition, fine-grained 15-state supervision and the Transformer-bridge also substantially improved gene model accuracy (Supplemental Figure 3). For the rest of this paper, we use the PlantGeneAnn 40-species model, hereafter referred to as PlantGeneAnn.

To demonstrate PlantGeneAnn’s predictive superiority, we next constructed a comprehensive benchmark and evaluated whole-genome *ab initio* annotation against three current SOTA tools (Helixer, Tiberius-v2.0.6, and ANNEVO-v2.3.3). The benchmark included 20 species: 18 angiosperms (including Arabidopsis and rice, both with the highest-quality reference annotations), one bryophyte (*Physcomitrium patens*), and one lycophyte (*Selaginella moellendorffii*) (Figure 1C). Notably, none of the benchmark species overlaps with the PlantGeneAnn training species, and comparisons were performed at the exon-, intron-, and locus-level (Supplemental Methods).

For the results, PlantGeneAnn significantly outperformed all three baseline tools at all three levels, establishing it as a SOTA in structural accuracy for plant gene annotation. Across the 20 species, mean F1 scores were 0.856 at the exon-level and 0.880 at the intron-level, exceeding ANNEVO, Tiberius, and Helixer by 1.7, 7.7, and 13.6 percentage points and by 1.2, 6.4, and 14.9 percentage points, respectively (Figure 1C; Supplemental Figure 4). Its advantage was greater in recovering complete gene structures: the mean locus-level F1 score reached 0.706, improving by 3.3 percentage points over ANNEVO and 13.6 and 22.5 percentage points over Tiberius and Helixer, respectively (Figure 1C, Supplemental Table 2). These results establish PlantGeneAnn as a more accurate annotator across the full spectrum of gene structure prediction, from local exon–intron boundary recognition to complete gene locus reconstruction.

We then performed more fine-grained precision–recall analyses. In Arabidopsis, rice, and *P. patens*, PlantGeneAnn achieved the highest precision among the four tools at both exon– and locus-level. Although its exon-level recall was slightly lower than that of ANNEVO in rice and *P. patens*, its locus-level recall exceeded that of ANNEVO in Arabidopsis and *P. patens* and was comparable in rice (Figure 1D). A species-grouped mean analysis showed a similar pattern: across the 18 angiosperms, PlantGeneAnn had significantly higher mean precision at both exon-and locus-level, whereas its mean recall was slightly lower than ANNEVO and higher than Tiberius and Helixer (Supplemental Figures 6–8). In the bryophyte and lycophyte, it retained higher precision and achieved the highest locus-level recall, indicating strong cross-lineage transferability (Figure 1E). BUSCO analysis further supported annotation completeness, with mean scores of 92.9% across all 20 species. ANNEVO attained a slightly higher mean BUSCO score (93.8%) (Supplemental Figure 5; Supplemental Table 3), consistent with its higher mean locus-level recall across the 20 species and its recall-oriented precision–recall trade-off, whereas PlantGeneAnn showed the strongest fine-grained structural accuracy. The pretrained GFM provides PlantGeneAnn with stronger contextual representations, enabling more accurate conversion of complex genomic sequences into gene models.

Having established PlantGeneAnn as an accurate *ab initio* annotation tool for plants, we next tested whether it could be adapted through fine-tuning to plant lineages outside angiosperms, such as the bryophyte *P. patens*, which performed poorly in the above benchmark. We applied LoRA fine-tuning and compared zero-shot and few-shot results, including Helixer, the only tool that provides an official interface for fine-tuning its released models, as a comparator. Fine-tuning substantially improved both tools: LoRA-fine-tuned PlantGeneAnn increased the locus-level F1 score from 0.581 to 0.736, a 15.4 percentage point gain (Figure 1F; Supplemental Figure 9), and achieved 82.1% BUSCO completeness, close to the 82.7% of the reference annotation. Full-parameter fine-tuning also improved Helixer, whereas fine-tuning only its final layer produced little improvement. In few-shot experiments, PlantGeneAnn retained a clear benefit with only 10% of the original fine-tuning dataset. Performance largely plateaued at 40% of the data, when exon– and locus-level F1 scores reached 0.896 and 0.707, respectively, and BUSCO completeness reached 81.8% (Figure 1F; Supplemental Table 4). These results show that lineage-specific fine-tuning can improve PlantGeneAnn performance even with limited data (40% for *P. patens*).

To assess computational requirements, we selected six representative species ranging from the 0.13 GB genome of Arabidopsis to the 3.98 GB genome of barley and recorded end-to-end annotation time and peak system memory usage. PlantGeneAnn contains 195M parameters, compared with 15M (Zhang *et al*., 2026), 8M (Gabriel et al., 2024), and 2M (Holst *et al*., 2026) for ANNEVO, Tiberius, and Helixer, respectively. Accordingly, PlantGeneAnn required more time than ANNEVO and Tiberius but remained faster than Helixer. However, its major advantage was low memory requirement: peak system memory usage of Helixer, ANNEVO, and Tiberius reached 69.6, 83.6, and 146.2 GB on the barley genome, whereas PlantGeneAnn never exceeded 18.8 GB on any of the six genomes, remaining well within the 32 GB of a typical personal workstation (Figure 1G; Supplemental Table 5). This efficiency lowers the hardware barrier for PlantGeneAnn and enables annotation of diverse plant genomes on standard personal workstations.

Finally, for complex application scenarios, we illustrate PlantGeneAnn’s performance on two complex gene loci: *OsYABBY4* in rice and *CAT3* in Arabidopsis. At the *OsYABBY4* locus (Ge et al., 2023), PlantGeneAnn reproduced the AGIS1.0 reference structure, whereas all three baseline tools predicted an additional exon between exons 5 and 6 (Figure 1H). At the *CAT3* locus (Zhang et al., 2012), PlantGeneAnn correctly recovered the complete gene structure, whereas Helixer incorrectly predicted the start codon, and ANNEVO and Tiberius misidentified the splice donor site downstream of exon 10 (Figure 1I). Together with the genome-wide benchmarking results, these examples indicate that PlantGeneAnn more accurately identifies exon–intron boundaries in complex gene loci.

## Conclusion

In summary, PlantGeneAnn is a plant-optimized, GFM-fine-tuned tool for accurate *ab initio* gene annotation. Without RNA-seq or homologous protein evidence, it predicts gene structures across diverse plant lineages, from exon boundaries to complete loci, particularly valuable for new assemblies and non-model plants. Its key innovation is using a GFM as the training starting point, significantly outperforming deep-learning baselines and establishing a SOTA in structural annotation accuracy. PlantGeneAnn can also be fine-tuned for specific lineages. Its greatest contribution is enabling accurate, efficient gene structure inference for large plant genomes on standard workstations, lowering the practical barrier. A current limitation is that PlantGeneAnn does not predict untranslated regions (UTRs); its output is restricted to complete protein-coding structures from start to stop codons. Future improvements will focus on extending PlantGeneAnn to annotate UTRs and alternative splicing isoforms.

## Data and code availability

The detailed training species, assembly versions, and training hyperparameters for PlantGeneAnn are provided in Supplemental Table 1. Detailed strategies for its fine-tuning and benchmarking are provided in Supplemental Methods. The code for the PlantGeneAnn annotation pipeline, together with its data-preprocessing, training, and fine-tuning pipelines, is available at https://github.com/qzzhang0131/PlantGeneAnn. The model weight for PlantGeneAnn is available at https://huggingface.co/qzzhang/PlantGeneAnn-v2-Angiospermae. The *ab initio* annotation GFF3 files generated by PlantGeneAnn for key model plants/crops are available at: https://huggingface.co/datasets/qzzhang/Key-plant-species-annotated-by-PlantGeneAnn.

## Author contributions

X.-H.H. and Q.-Y.Y. designed, supervised the research and revised the manuscript. Q.-Z.Z., and Z.-Y.Z. collected the data and built the PlantGeneAnn model. Q.-Z.Z. designed and wrote the PlantGeneAnn annotation pipeline. Q.-Z.Z., Z.-Y.Z., K.-P.L., and J.W. performed the gene annotation benchmarks. Q.-Z.Z. and Z.-Y.Z. performed the ablation analysis. Q.-Z.Z. performed the fine-tuning study and few-shot learning. Z.-Y.Z. and K.-X.D. analyzed annotations at two loci. Q.-Z.Z. and Z.-Y.Z. wrote the manuscript. All authors read and approved the final manuscript.

## Supplemental Figures

**Supplemental Figure 1.**
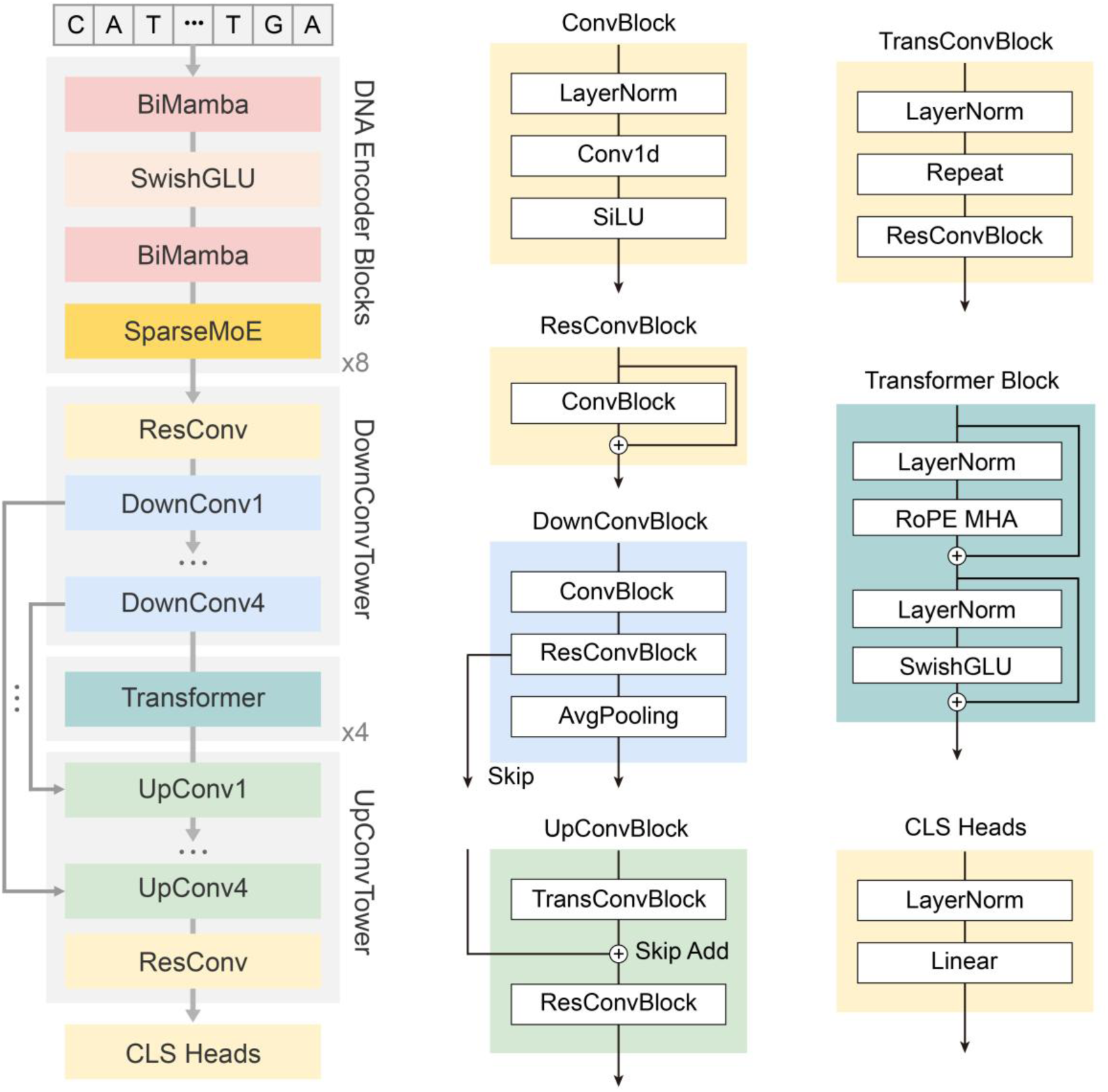
PlantGeneAnn model architecture. PlantGeneAnn model consists of four sequential components: a DNA encoder, a down-convolution tower, a transformer bridge and an up-convolution tower.

**Supplemental Figure 2.**
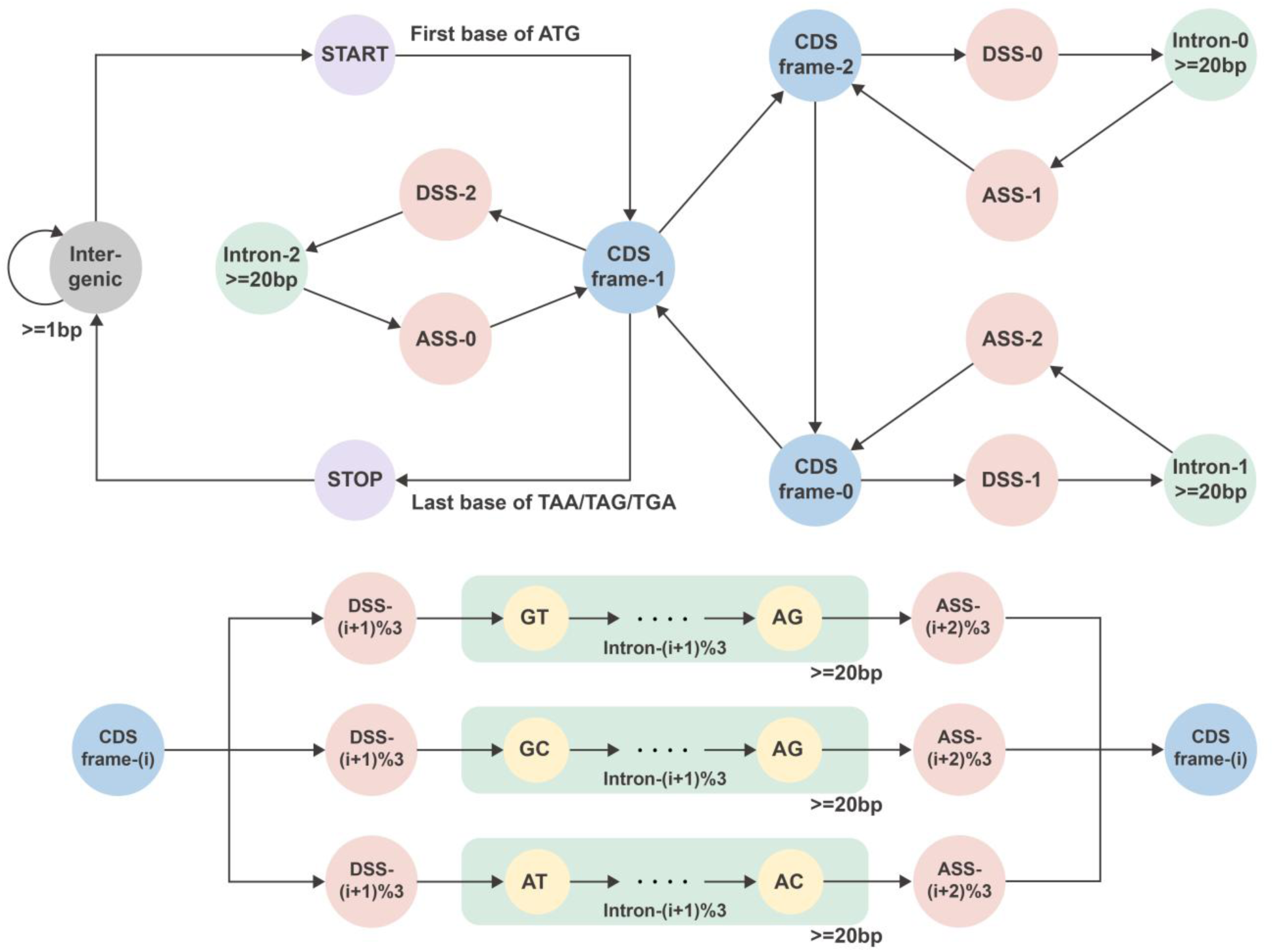
Biological grammar of the PlantGeneAnn semi-Markov model (SMM) decoder. Arrows indicate the allowed state transitions in the transcriptional direction. A gene enters the three-frame CDS cycle from the START state at the first base of the ATG start codon and terminates through the STOP state at the last base of the in-frame TAA, TAG, or TGA stop codon; phase-coupled DSS–intron– ASS transitions maintain reading-frame continuity across exons. Introns are at least 20 bp in length, and only the GT–AG, GC–AG, and AT–AC splice-site combinations are permitted. The lower panel summarizes the intron transition rules for an arbitrary CDS frame-i.

**Supplemental Figure 3.**
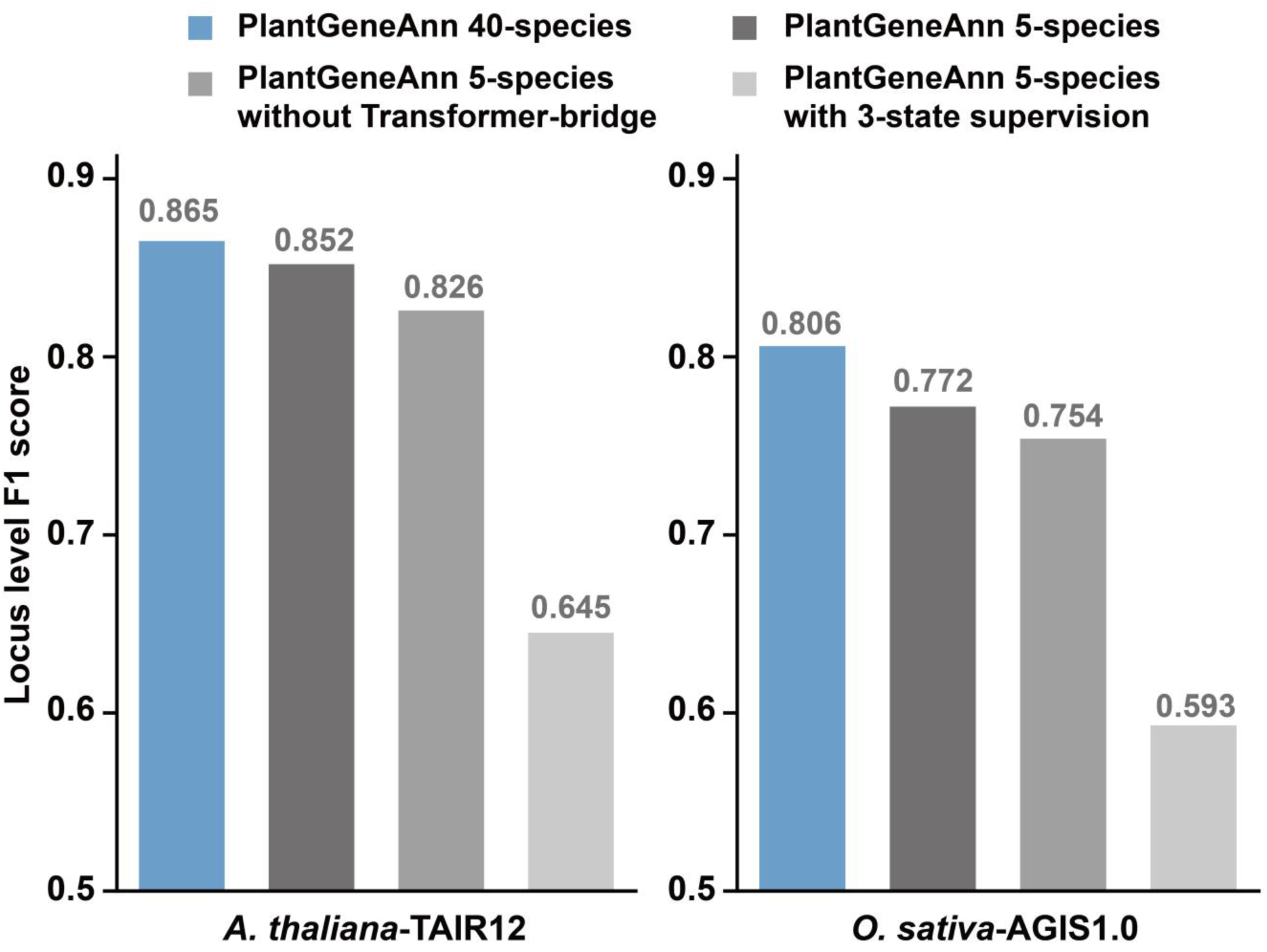
Ablation analysis of PlantGeneAnn. Locus-level F1 scores for PlantGeneAnn and its ablated variants evaluated on whole-genome annotations of Arabidopsis (TAIR12) and rice (AGIS1.0). Bars compare the 40-species PlantGeneAnn model, a five-species model, a five-species model without the Transformer bridge, and a five-species model with the output states reduced from 15 to 3 states (protein-coding gene, exon, CDS). Values above the bars indicate the corresponding locus-level F1 scores.

**Supplemental Figure 4.**
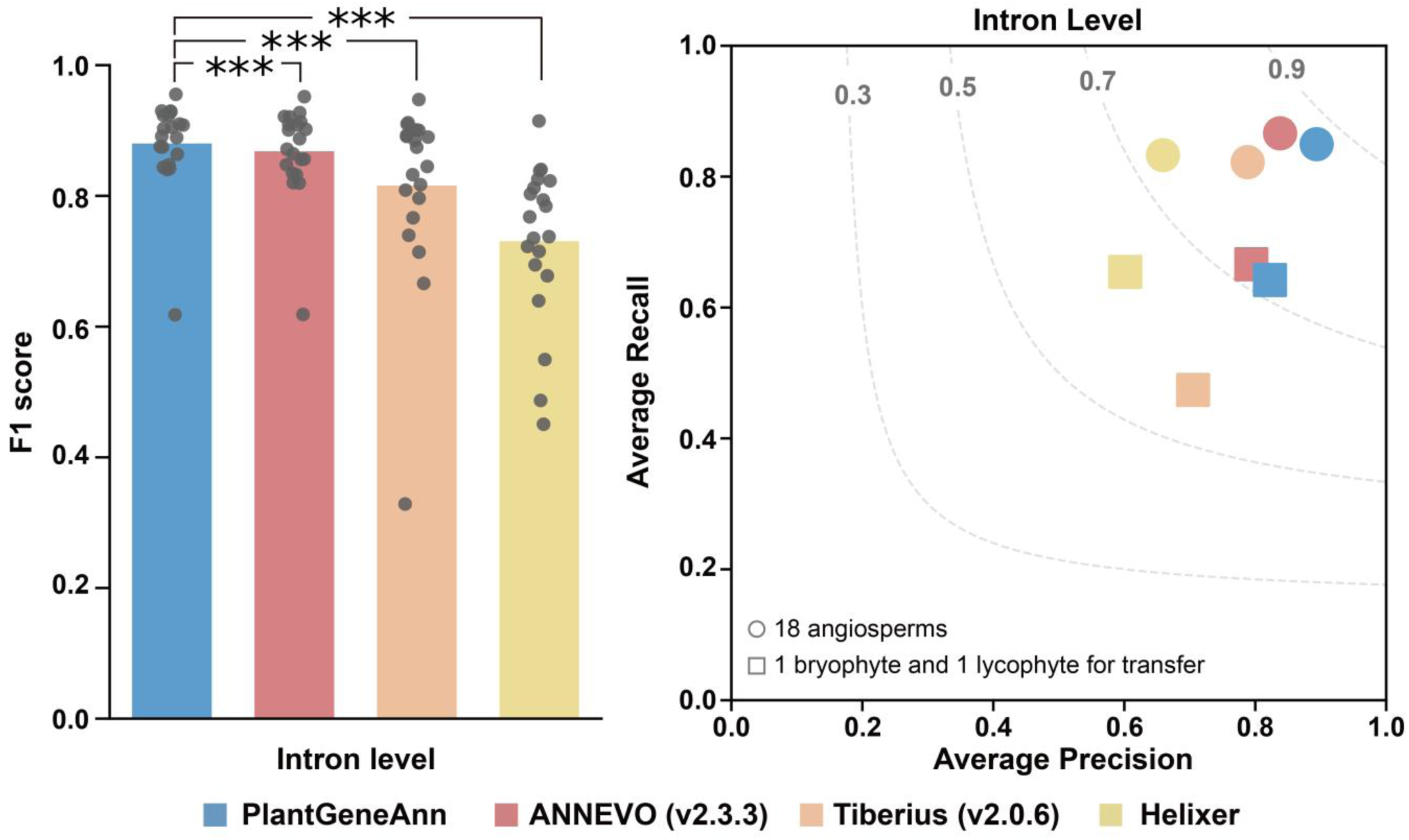
Intron-level accuracy comparison across 20 test species. Left, intron-level F1 scores for the four annotation tools; bars indicate the mean score across species, and points represent species-specific scores. Asterisks indicate significant differences of PlantGeneAnn versus each baseline tool (two-sided paired Wilcoxon signed-rank tests; *** P < 0.001). Right, average precision– recall performance at the intron level. Circles represent the 18 angiosperm species, whereas squares represent one bryophyte and one lycophyte included to assess cross-clade transfer. Dashed contours indicate intron-level F1 scores.

**Supplemental Figure 5.**
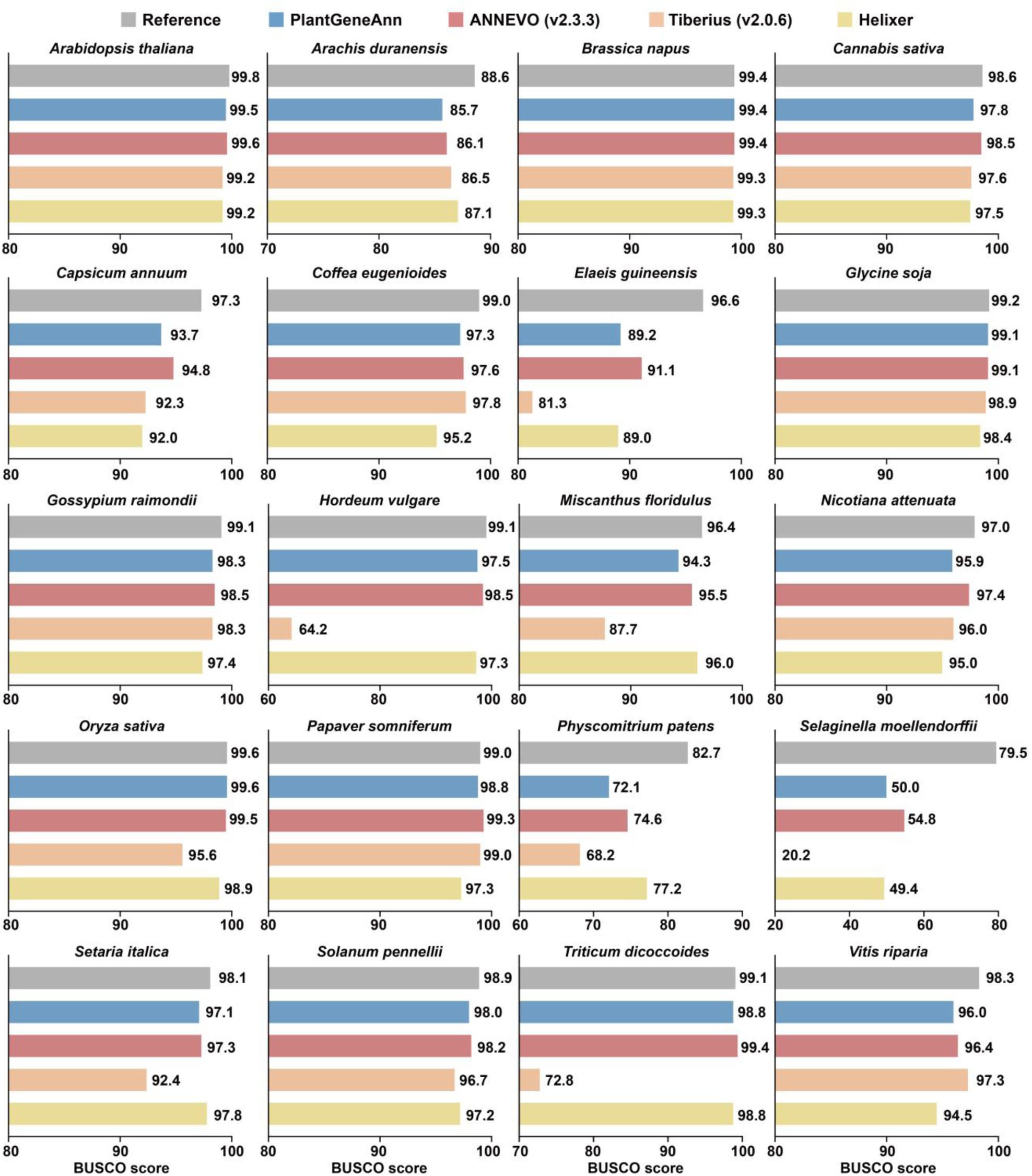
BUSCO completeness comparison across 20 test species. BUSCO completeness scores for the reference annotations and predictions generated by PlantGeneAnn, ANNEVO, Tiberius, and Helixer across 20 test species. Horizontal bars show the BUSCO score for each annotation set in each species.

**Supplemental Figure 6.**
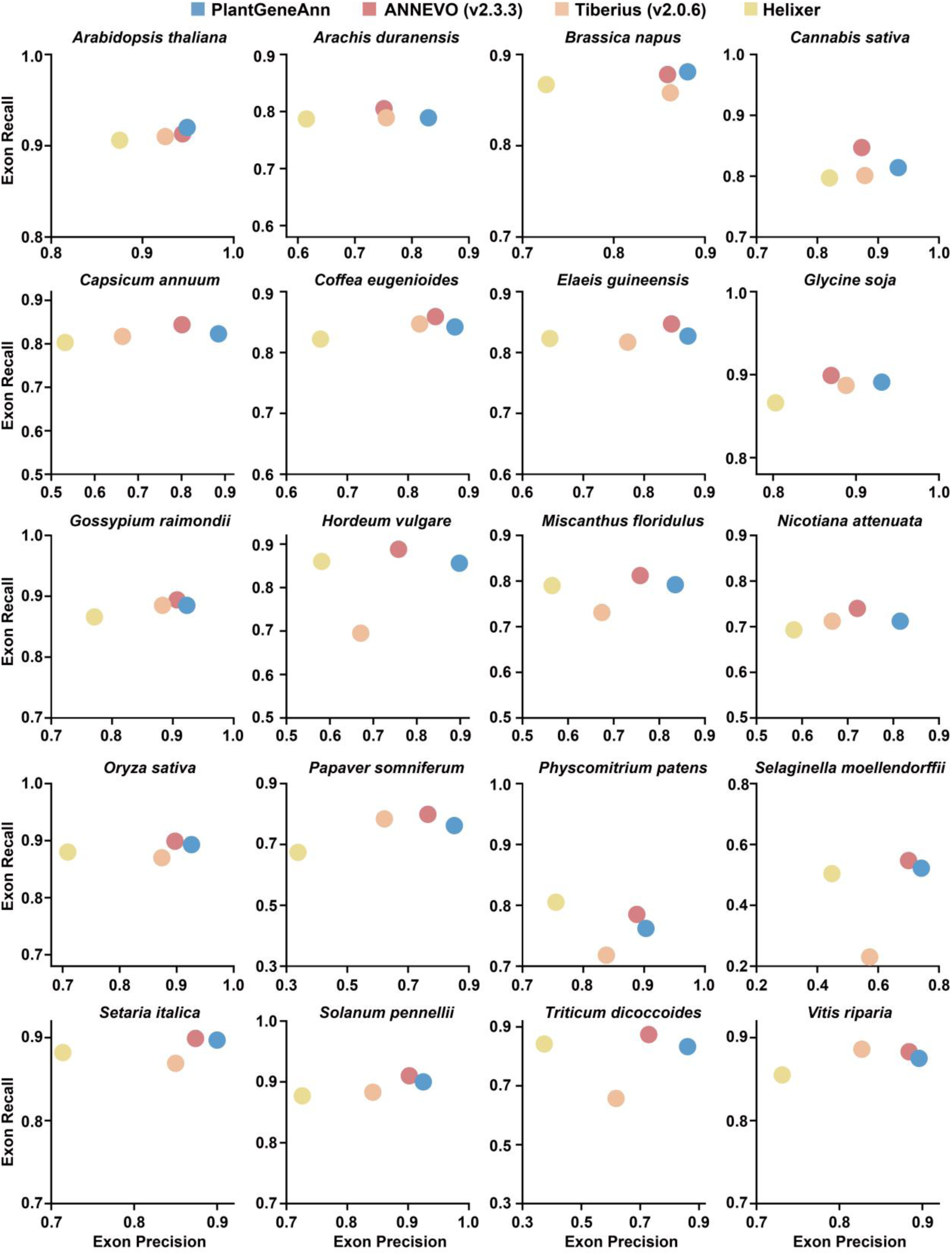
Exon-level precision–recall analysis across 20 test species.

**Supplemental Figure 7.**
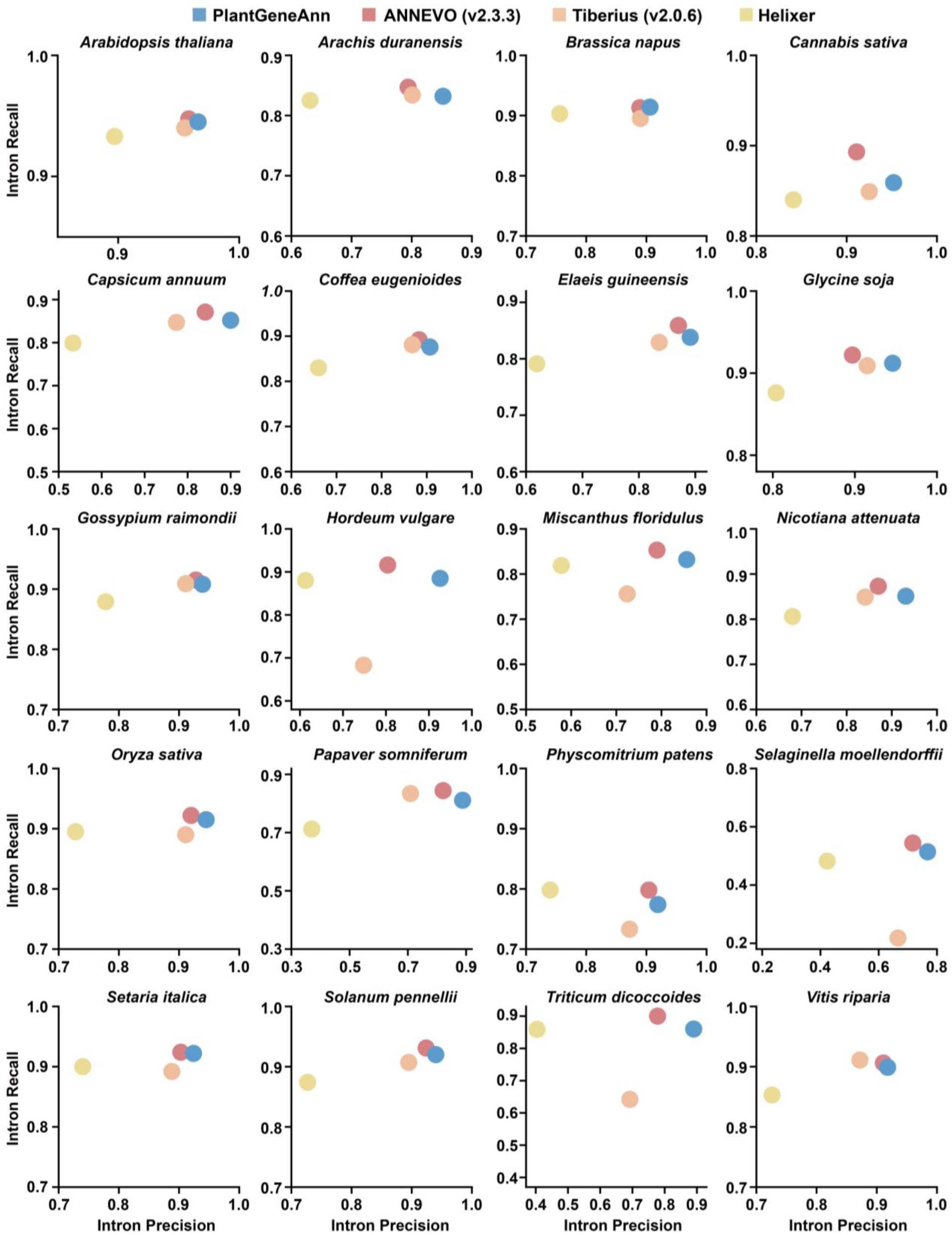
Intron-level precision–recall analysis across 20 test species.

**Supplemental Figure 8.**
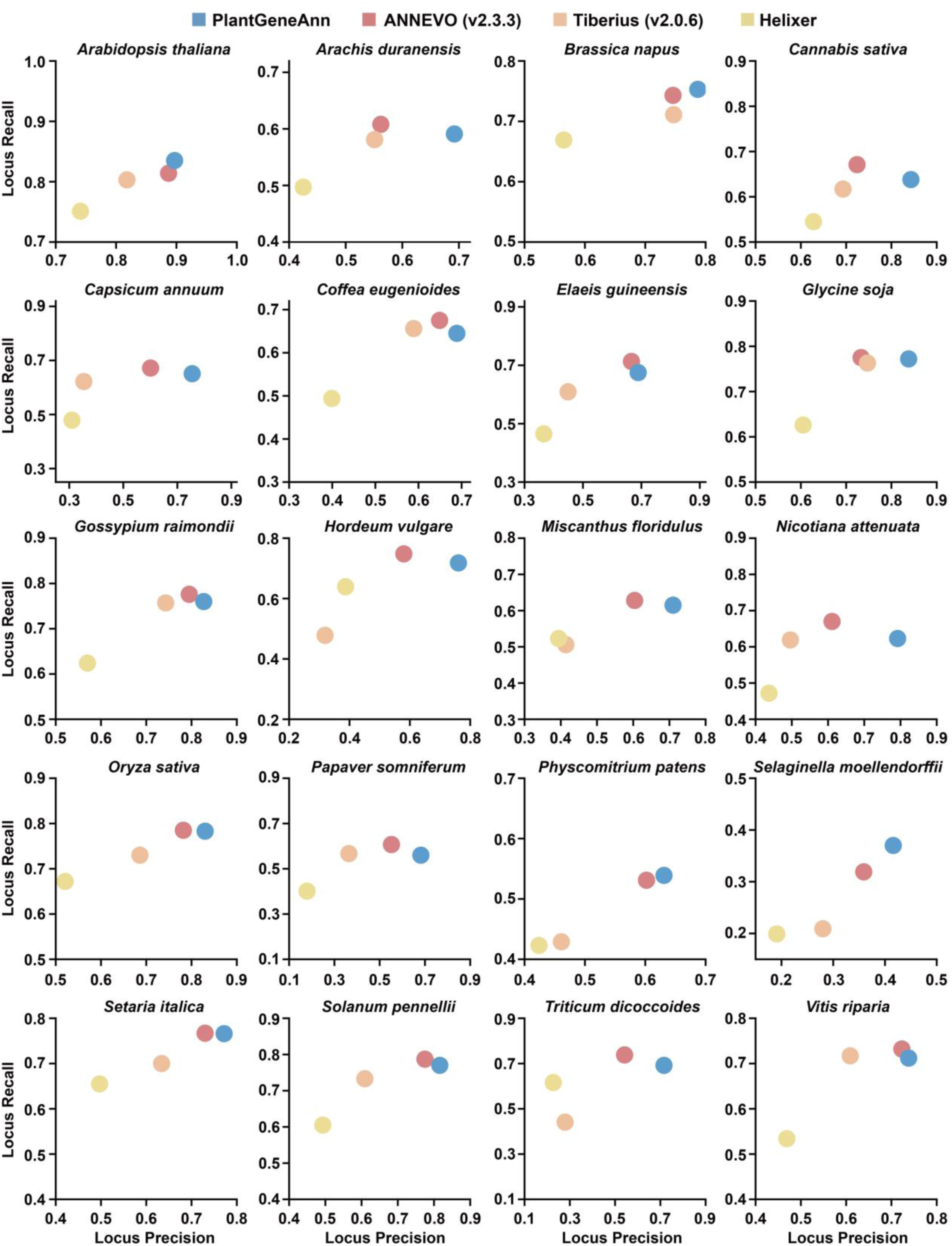
Locus-level precision–recall analysis across 20 test species.

**Supplemental Figure 9.**
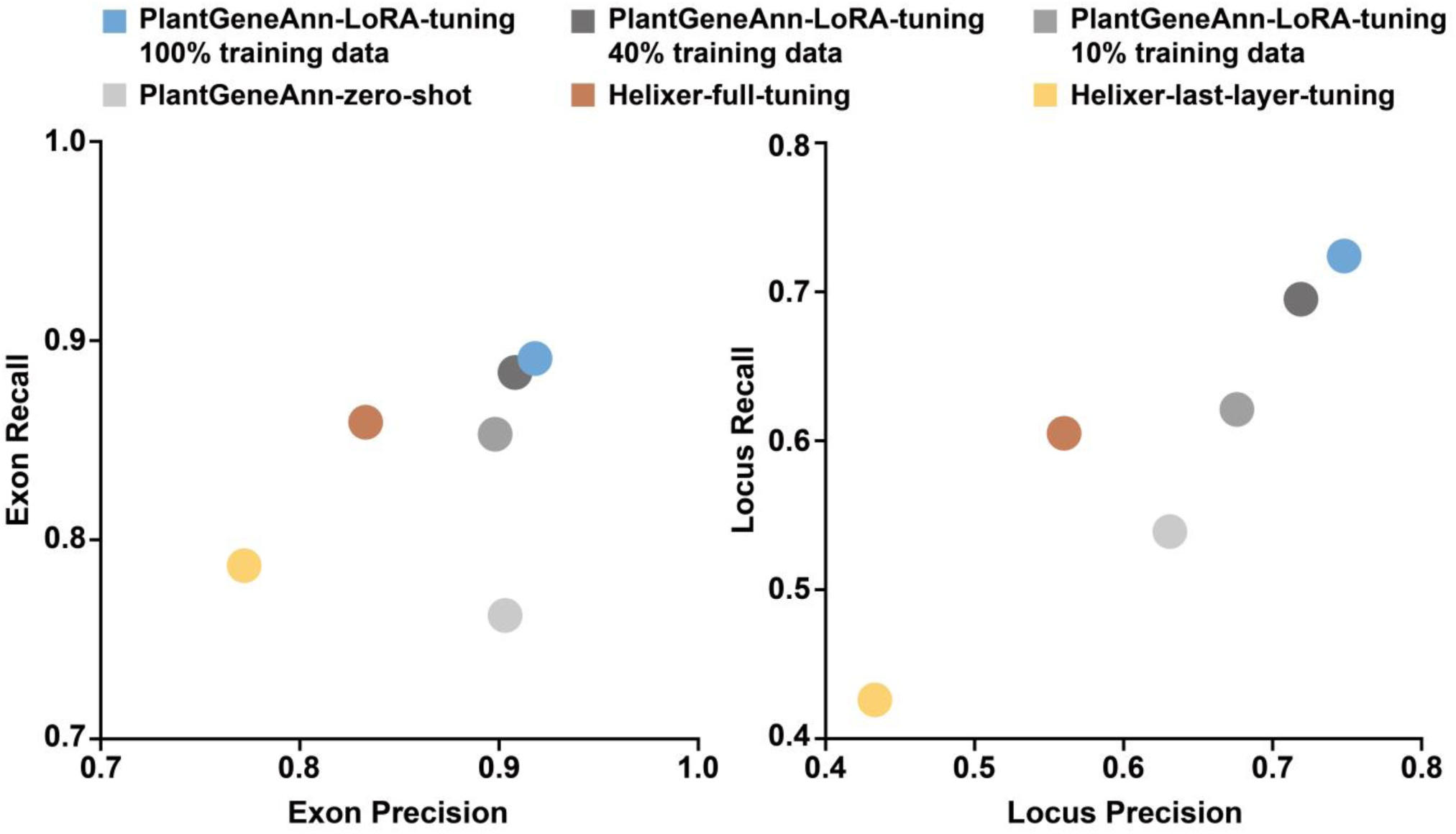
Exon– and locus-level precision–recall analyses of fine-tuning performance on *Physcomitrium patens*. The left panel plots exon-precision against exon-recall, and the right panel plots locus-precision against locus-recall. Points correspond to PlantGeneAnn with LoRA fine-tuning using 100%, 40%, or 10% of the fine-tuning data, PlantGeneAnn in the zero-shot setting, Helixer with full-parameter fine-tuning, and Helixer with last-layer fine-tuning.

## Supplemental Methods

### PlantGeneAnn model architecture

PlantGeneAnn is a base-resolution sequence segmentation model designed for *ab initio* annotation of plant genomes. The model consists of a DNA encoder and an embedding decoder connected in series. The DNA encoder is composed of 16 bidirectional Mamba blocks (Schiff et al., 2024), with layers alternately employing SwishGLU (Touvron et al., 2023) and sparse mixture-of-experts (SparseMoE) modules (Shazeer et al., 2017), and is responsible for extracting high-resolution single-nucleotide embedding representations from raw DNA sequences. The embedding decoder is built upon a one-dimensional U-Net-style architecture, comprising a 4-level down-sampling convolutional tower and a 4-level up-sampling convolutional tower (Ronneberger et al., 2015). To capture long-range dependencies in genomic sequences, we introduce a 4-layer Transformer encoder (Vaswani et al., 2017) with rotary position embeddings (RoPE) (Su et al., 2024) at the bridge between the down-sampling and up-sampling towers. The final classification head consists of a linear mapping that projects the 512-dimensional representation at each position into 30 logits. The first 15 channels represent 15 mutually exclusive gene structure states on the forward strand, and the last 15 channels represent the corresponding states on the reverse strand. The set of states on each strand is adapted from the Tiberius state set (Gabriel et al., 2024) and includes: intergenic region, three intron phase states, three CDS phase states, an open reading frame (ORF) start site state (START), three donor splice site (DSS) states, three acceptor splice site (ASS) states, and an ORF termination state (STOP). The full model contains approximately 195 million trainable parameters.

### Semi-Markov model architecture

To convert the per-base state probabilities produced by the neural network into structurally complete and coding-grammar-compliant gene models, we designed a strict 15-state semi-Markov model (SMM) decoder. Decoding proceeds independently along the transcriptional direction of the forward and reverse strands. For each position, the 15-class softmax probabilities from the model are first transformed into calibrated log-odds scores relative to the intergenic state, serving as state emission scores:

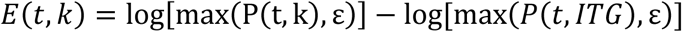

where *P*(*t*, *k*) denotes the predicted probability that position *t* belongs to state *k*, *P*(*t*, *ITG*) denotes the probability of the intergenic state, and *ε* is a small constant to avoid taking the logarithm of zero. The score of an all-intergenic path is defined as zero, allowing the decoder to return an empty prediction when insufficient gene evidence is present.

A valid protein-coding gene must start with an ATG initiation codon and terminate with TAA, TAG, or TGA in the same reading frame. CDS states advance in a fixed cyclic order of frame-0, frame-1, and frame-2, and in-frame stop codons are prohibited within a complete CDS. The full CDS length is constrained to be at least 60 bp, and its length is guaranteed to be a multiple of 3 by the state grammar (Holst et al., 2026). Unlike the per-base hidden Markov models (HMMs) classically used for gene finding (Lukashin and Borodovsky, 1998) and employed by annotation tools such as ANNEVO (Zhang et al., 2026) and Tiberius (Gabriel et al., 2026), our method represents introns as variable-length segment edges from a donor site to an acceptor site. For a donor splice site state with phase i (DSS-i), a valid path sequentially traverses DSS-i, intron state intron-i, and acceptor splice site state ASS-((i+1) mod 3), before resuming the corresponding CDS phase. Only three classes of splice site combinations–GT-AG, GC-AG, and AT-AC–are permitted, and the intron length must be no shorter than 20 bp. For introns that interrupt a codon (i.e., split codons), the algorithm retains partial codon context in the vicinity of the donor site and reconstructs the complete triplet codon after splicing at the acceptor site; any candidate splice junction that introduces a premature stop codon is excluded. To avoid enumerating all donor–acceptor pairs, we construct prefix sums of the emission scores for each of the three intron phases and group donor candidates that satisfy the minimum length constraint by phase, donor motif, split-codon context, and truncated CDS length. Each group retains only the highest-scoring donor candidate, enabling constant-time calculation of the cumulative score for a complete intron segment upon scanning an acceptor site.

The dynamic programming (DP) simultaneously maintains path backtracking information and a list of completed genes, jointly optimizing the total score of zero, one, or multiple non-overlapping complete genes within each candidate region. Finally, exon intervals in the transcriptional direction are recovered via backtracking, and results on the reverse strand are remapped to the original genome coordinates.

### Construction of training and validation datasets

To train PlantGeneAnn, we constructed a training set comprising 40 angiosperm species from diverse evolutionary clades and a validation set comprising 8 species (all training and validation species are listed in Supplemental Table 1). For each species, genome FASTA files and corresponding GFF3 annotations were downloaded from the NCBI RefSeq database (accessed June 15, 2026) (O’Leary et al., 2016), and GFF annotation files were processed in the following two steps:

(1) All protein-coding genes were extracted from the GFF annotation files, and the primary transcript of each gene (defined as the transcript encoding the longest protein sequence) and its associated annotation entries were retained.
(2) All genes retained in the previous step were subjected to quality control, during which non-canonical ORFs and low-confidence gene models were filtered out (including non-canonical start codons, non-canonical stop codons, non-canonical splice junctions, premature termination, partial genes, introns shorter than 20 bp, and ambiguous bases in coding regions or splice sites), and the GeneIDs of all filtered genes were recorded.

For all training genome files, we segmented the genomes into equal-length fragments using a window length of 40,960 bp and a step size of 30,720 bp, and removed fragments with more than 1% ambiguous bases. We then generated 15-state labels for each of the forward and reverse strands following the Tiberius training pipeline (Gabriel *et al*., 2026), and used the filtered gene regions to generate a loss mask. The detailed data preprocessing pipeline is available at: https://github.com/qzzhang0131/PlantGeneAnn.

For all validation genome files, we segmented the genomes into non-overlapping 40,960 bp fragments, removed fragments with more than 1% ambiguous bases, and generated only the 15-state labels for each of the forward and reverse strands.

### PlantGeneAnn training

When training PlantGeneAnn, the DNA encoder was initialized with our previously published genome foundation model PlantBiMoE (https://huggingface.co/qzzhang/PlantBiMoE-116M-copy) to leverage its prior knowledge (Lin et al., 2025). We optimized the model using the AdamW optimizer (Loshchilov and Hutter, 2017), with β1 and β2 set to 0.9 and 0.99, respectively, a weight decay of 0.01, and an effective batch size of 32 (achieved via gradient accumulation across multiple GPUs). To mitigate the decrease in prediction accuracy at sequence boundaries caused by asymmetric visibility, we computed the loss and backpropagated gradients only over the central 80% of the input sequence (i.e., the central 32,768 bp region); this strategy has been widely adopted in previous related studies (Linder et al., 2025) (Zhang *et al*., 2026). In terms of training step configuration, we adopted 10,000 steps as one unit (corresponding to 13.1 billion tokens) and set the maximum number of training steps to 400,000. The learning rate was linearly warmed up from 0 to 1e-4 over the first 5% of training steps, followed by linear decay to 5e-5 over the remaining steps. During training, model checkpoints were saved and validation was performed every 10,000 steps. We used the macro-averaged F1 score (macro-F1) across the 30 class labels on both strands in the validation set as the monitoring metric, and early stopping was triggered when this metric failed to improve.

### Hierarchical masked Focal–F1 loss

To effectively mitigate the extreme class imbalance commonly encountered in base-resolution gene structure annotation–where intergenic bases vastly outnumber functional states–we designed a hierarchical masked Focal–F1 loss. This loss function comprises four weighted terms on a single strand: a masked focal classification loss (Lin et al., 2017), a non-intergenic state-level soft-F1 loss (Gabriel *et al*., 2024), an intron-group soft-F1 loss, and a coding-group soft-F1 loss, formulated as:

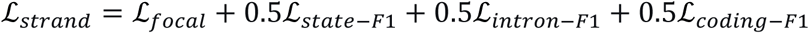

#### Masked focal classification

Let *p_t_* denote the predicted probability of the true class at position *t*, *m_t_* ∈ {0, 0.5, 1} the loss mask, and *α_yt_* the corresponding class weight. The masked focal loss is defined as:

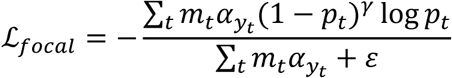

where the focusing parameter *γ* = 2 and *ε* is a numerical stability term. In the class weight vector, all 14 states are assigned a weight of 1, except for the start codon state (START, index 7), which is assigned a weight of 2 to moderately emphasize accurate prediction of the start site.

#### Hierarchical soft-F1 loss

For any state or state group *g*, we adopt a soft-F1-style loss:

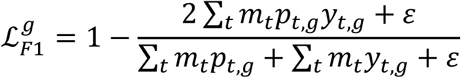

where *p_t_*_,*g*_ and *y_t_*_,*g*_ denote the predicted probability and the ground-truth label indicator for that group, respectively. The state-level term ℒ_state_ covers all 14 non-intergenic states (indices 1–14). The intron-group term ℒ_intron−F1_ is obtained by merging states 1–3 (corresponding to the three intron-related states). The coding-group term ℒ_coding−F1_ is obtained by merging states 4–14, encompassing the three CDS reading frames and their start, stop, and six splice sites (ASS, DSS) states.

When a state or state group is completely absent from a training sample, the above soft-F1 loss reduces to a false-positive penalty term based on the predicted probabilities, thereby effectively suppressing spurious activation of missing classes. For the state-level missing penalty, we further introduce a softmax gating mechanism with a temperature parameter *τ* = 0.20 to obtain more stable gradient signals.

#### Joint optimization over both strands

The strand loss ℒ_strand_ is computed independently for the forward and reverse strands, and their arithmetic mean is taken as the final training objective:

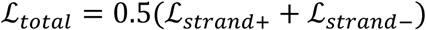

This hierarchical design maintains the focus of the focal loss on hard examples while explicitly optimizing the recall and precision of functional states through multi-granularity soft-F1 terms, thereby improving the prediction of complex gene structures, especially multi-exon genes and boundary sites.

### Ablation analysis

To assess the impact of key components in PlantGeneAnn, we conducted ablation studies. In addition to the complete PlantGeneAnn model trained on 40 angiosperm species, we selected 5 species from this training set and trained a variant, designated PlantGeneAnn-5-species, using exactly the same strategy, and used it as the baseline model for ablation. Subsequently, while keeping all other components unchanged, we performed the following replacements: (1) directly converting the raw model predictions into gene models without SMM decoding (without SMM); (2) replacing the GFM-initialized weights with randomly initialized weights (without GFM-init); (3) replacing the Transformer bridge in the embedding decoder with a convolution bridge (without Transformer-bridge); (4) replacing the fine-grained 15-state supervision combined with the masked focal-F1 loss with 3-state (protein-coding gene, exon, CDS) supervision combined with the weighted binary cross-entropy (BCE) loss (3-state supervision). All four ablated models were trained using the same training procedure and training data as the baseline model PlantGeneAnn-5-species. Finally, we selected the converged weights of all models and compared their whole-genome annotation results on two model plants from the main benchmark, *Arabidopsis thaliana* (TAIR12) and *Oryza sativa* (AGIS1.0), neither of which belongs to any ablation training set. We reported the most important locus-level F1 scores.

### Construction of the test set

To comprehensively evaluate the performance of annotation models, we constructed a test set comprising 20 plant species from diverse evolutionary clades. First, following the practice of ANNEVO v2.3.3 (Zhang *et al*., 2026), we included *Arabidopsis thaliana* (TAIR12) and *Oryza sativa* (AGIS1.0)–two botanical “gold standards”–in the test set. Second, we selected seven species from the Helixer test set (Holst *et al*., 2026) (*Brassica napus*, *Cannabis sativa*, *Nicotiana attenuata*, *Solanum pennellii*, *Papaver somniferum*, *Triticum dicoccoides*, *Vitis riparia*) and nine species from NCBI RefSeq (*Arachis duranensis*, *Capsicum annuum*, *Coffea eugenioides*, *Elaeis guineensis*, *Glycine soja*, *Gossypium raimondii*, *Hordeum vulgare*, *Miscanthus floridulus*, *Setaria italica*) as supplements. Together, these formed a test set of 18 angiosperm species, including 7 rosids, 4 asterids, 6 monocots, and one basal eudicot (Papaveraceae, Ranunculales). Finally, we also included *Physcomitrium patens* (a bryophyte) and *Selaginella moellendorffii* (a lycophyte) to assess the transfer learning capability of each model. The genome assemblies of all test species were updated to the latest versions available as of June 15, 2026 (exact NCBI RefSeq accession numbers for all genomes and corresponding annotations are provided in Supplemental Table 1), and none of these species were present in the PlantGeneAnn training set. It should be noted that, because we could not fully align the training data of all annotation models included in the comparison, some test species may appear in the training data of certain comparative models. Nevertheless, we prioritized the comprehensiveness, representativeness, and annotation quality of the test species when constructing the test set.

### Benchmarking against existing *ab initio* annotation tools

To minimize confounding variables, all *ab initio* annotations were performed on the same server equipped with two NVIDIA RTX 4090 GPUs and 36 vCPUs of an AMD EPYC 9754 processor. For a given genome FASTA file `genome.fa` and the pretrained model `PlantGeneAnn-v2-Angiospermae`, PlantGeneAnn v2.0 was run as follows:

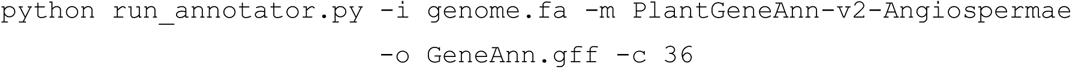

#### Comparison with ANNEVO

ANNEVO v2.3.3 (Zhang *et al*., 2026) was used for *ab initio* annotation of all genomes. For a given genome FASTA file `genome.fa`, the following command was used:

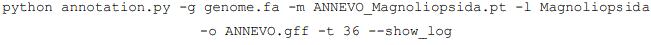

The Magnoliopsida model is the only plant model provided by ANNEVO; following the NCBI Taxonomy (TaxID: 3398), it covers all flowering plants, including both monocots and dicots, and is therefore the recommended model for the angiosperm test species.

#### Comparison with Tiberius

Tiberius v2.0.6 (Gabriel *et al*., 2026) was used for *ab initio* annotation of all genomes. For a given genome FASTA file `genome.fa`, the following command was used:

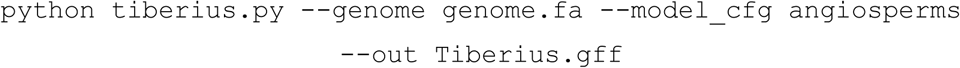

#### Comparison with Helixer

Helixer v0.3.6 (Holst *et al*., 2026) was used for *ab initio* annotation of all genomes. For a given `species` with genome FASTA file `genome.fa`, the following command was used:

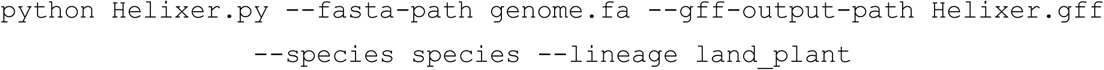

### Fine-tuning study and few-shot learning on *Physcomitrium patens*

#### Fine-tuning dataset

To enable the PlantGeneAnn model weights to learn gene structures of bryophytes, we downloaded the genome and annotation files of *Ceratodon purpureus* and *Sphagnum fallax* from the Phytozome v13 database (accessed June 15, 2026) (Goodstein et al., 2012), and used *Marchantia polymorpha* as the validation set for training monitoring. All genome and annotation files were processed uniformly using the PlantGeneAnn data preprocessing pipeline described above.

#### Fine-tuning study

To efficiently fine-tune PlantGeneAnn while mitigating catastrophic forgetting to some extent, we applied LoRA for parameter-efficient fine-tuning (PEFT) of PlantGeneAnn (Hu et al., 2021). Specifically, LoRA adapters were injected into the input projection layers (`in_proj`) and output projection layers (`out_proj`) of the 16 BiMamba blocks in the DNA encoder, while all other parameters remained frozen. For the embedding decoder, we injected LoRA into all linear projection layers in the transformer bridge, and unfroze only the parameters of the last up-convolution (UpConv) block, the local-refine residual-convolution (ResConv) block, the final normalization layer, and the prediction head, while keeping the remaining modules frozen. All LoRA injections used a rank of 8 and an alpha of 8. Overall, this scheme introduced learnable parameters accounting for approximately 2.45% of the total PlantGeneAnn parameters (approximately 4.8M). Fine-tuning was also performed using the AdamW optimizer (Loshchilov and Hutter, 2017), with β1 and β2 set to 0.9 and 0.99, respectively, a weight decay of 0.01, and an effective batch size of 32. We set 20 epochs; during training, model checkpoints were saved and validation was performed after every epoch. The learning rate was linearly warmed up from 0 to 1e-4 during the first epoch, and then decayed to 0 following a cosine schedule over the remaining epochs. We used the macro-F1 across the 30 class labels on both strands in the validation set as the monitoring metric, and early stopping was triggered when this metric failed to improve. Finally, we evaluated the fine-tuned model (https://huggingface.co/qzzhang/PlantGeneAnn-v2-Bryophyta) on *Physcomitrium patens* to obtain performance metrics.

#### Few-shot learning

For the fine-tuning dataset described above, we progressively reduced the number of samples to 80%, 60%, 40%, 20%, and 10% of the original size (each reduction was performed on the dataset obtained from the previous step; for example, the 60% dataset was derived by further subsampling the 80% dataset), yielding five fine-tuning datasets of different sizes. We trained five models to convergence using the fine-tuning strategy described above and evaluated their performance metrics on *Physcomitrium patens*.

#### Comparison with Helixer fine-tuning

We used two fine-tuning strategies provided in the Helixer GitHub repository (https://github.com/usadellab/Helixer)–full-parameter fine-tuning and last-layer fine-tuning–to fine-tune the Helixer `land_plant` model using the two bryophytes mentioned above as the training set, and similarly used *Marchantia polymorpha* as the validation set to monitor training. For ANNEVO and Tiberius, because their repositories do not provide an official interface for fine-tuning the released model weights to adapt to new lineages, and because training them from scratch on the two bryophyte genomes would be inconsistent with a fine-tuning strategy, we did not include them in this comparison.

### Annotation metrics

**Structural accuracy metrics**: the structural accuracy metrics were calculated largely following the evaluation protocol used by ANNEVO and Tiberius. First, all protein-coding genes were extracted from the reference annotation file, and entries containing non-canonical open reading frames (ORFs) were filtered out. Because none of PlantGeneAnn, ANNEVO, or Tiberius can predict UTRs, we removed UTRs from all transcripts before scoring. For the prediction file `query.gff`, gffcompare v0.12.10 (Pertea and Pertea, 2020) was first used to compare it against the filtered reference annotation `reference.gff` with the following command:

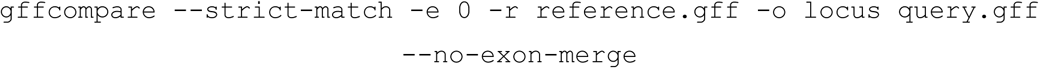

This yielded locus-level sensitivity and precision. Next, the primary (or longest) protein-coding transcript for each gene was extracted from `reference.gff` to generate `primary_isoform.gff`, and the following command was used for comparison:

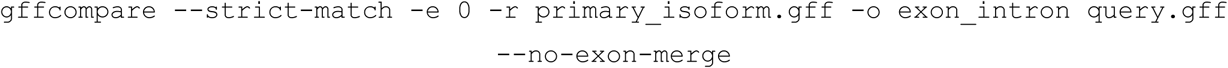

This yielded exon-level and intron-level sensitivity and precision. For all sensitivity (SN) and precision (PR) values, the F1 score was calculated as:

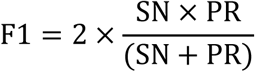

**BUSCO completeness:** as a complement to the structural accuracy metrics, we used BUSCO v6.0.0 (Tegenfeldt et al., 2025) to assess the completeness of all annotation files, including the reference annotation files. First, we used gffread v0.12.7 (Pertea and Pertea, 2020) to extract protein sequences into `query_prot.faa` from each annotation file `query.gff` using the following command:

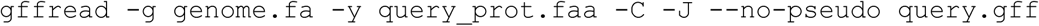

We then performed BUSCO analysis in protein mode using the following command:

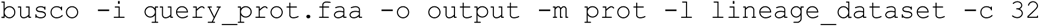

The lineage_dataset used for each species is provided in Supplemental Table 3.

### Runtime and peak memory usage comparison

To compare the runtime and peak system memory usage of different annotation tools, we performed *ab initio* annotation of six representative plant genomes of varying sizes–*Arabidopsis thaliana* (138 MB),

*Oryza sativa* (373 MB), *Cannabis sativa* (744 MB), *Zea mays* (2.06 GB), *Papaver somniferum* (2.57 GB), and *Hordeum vulgare* (3.98 GB)–using PlantGeneAnn, ANNEVO, Tiberius, and Helixer on a server equipped with a single NVIDIA RTX 4090 GPU and 18 vCPUs of an AMD EPYC 9754 processor, a configuration accessible to most laboratories or individual researchers. The accuracy benchmark and the runtime benchmark were conducted independently, each with the configuration stated in its respective section. For each tool, the batch size was set to the maximum value that could fit within GPU memory to maximize GPU utilization; for tools supporting multi-threaded parallel decoding, the number of decoding threads was uniformly set to 18 to maximize CPU utilization. The complete command-line parameters for each tool on each genome are provided in the section “Benchmarking against existing *ab initio* annotation tools” above. We recorded the wall-clock time from job submission to annotation file output, as well as the peak system memory usage during this period. For each tool and each genome, annotations were repeated twice, and the average runtime and average peak memory usage were used.

### Hardware

PlantGeneAnn was trained on a server equipped with eight NVIDIA A800 80GB GPUs, and the training process took approximately 80 hours to converge. Fine-tuning was performed on a server equipped with eight NVIDIA RTX 4090 GPUs and took approximately 3.5 hours to converge.

## References

1. Boshar, S., Evans, B., Tang, Z., Picard, A., Adel, Y., Lorbeer, F.K., Rajesh, C., Karch, T., Sidbon, S., Emms, D., et al. (2025). A foundational model for joint sequence-function multi-species modeling at scale for long-range genomic prediction. bioRxiv 10.64898/2025.12.22.695963.

2. Gabriel, L., Becker, F., Hoff, K.J., and Stanke, M. (2024). Tiberius: end-to-end deep learning with an HMM for gene prediction. Bioinformatics 40:btae685.

3. Gabriel, L., Brůna, T., Kaur, A., Krishnan, A., Ortmann, F., Salamov, A., Talbot, S., Becker, F., Krieg, R., and Wheat, C.W. (2026). Accurate ab initio gene prediction in eukaryotes with Tiberius in multiple clades. bioRxiv.

4. Ge, R., Ye, C., Peng, Y., Dai, Q., Zhao, Y., Liu, S., Wang, P., Hu, L., and He, C. (2023). m(6)A-SAC-seq for quantitative whole transcriptome m(6)A profiling. Nat Protoc 18:626–657. 10.1038/s41596-022-00765-9.

5. Holst, F., Bolger, A.M., Kindel, F., Günther, C., Maß, J., Triesch, S., Kiel, N., Saadat, N., Ebenhöh, O., Usadel, B., et al. (2026). Helixer: ab initio prediction of primary eukaryotic gene models combining deep learning and a hidden Markov model. Nature Methods 23:732–739. 10.1038/s41592-025-02939-1.

6. Ji, H.J., Pertea, M., and Salzberg, S.L. (2026). Annotating genomes at increased scale and resolution. Nature Reviews Genetics:1–13.

7. Lin, K., Zhang, Q., Wang, R., Hu, X., and Xu, W. (2025). PlantBiMoE: a bidirectional foundation model with SparseMoE for plant genomes. In 2025 IEEE International Conference on Bioinformatics and Biomedicine (BIBM). 10.1109/BIBM66473.2025.11356776.

8. Michael, T.P. (2026). Plant genome assembly and annotation. Current Opinion in Plant Biology 90:102859.

9. Stanke, M., and Waack, S. (2003). Gene prediction with a hidden Markov model and a new intron submodel. Bioinformatics 19:ii215–ii225. 10.1093/bioinformatics/btg1080.

10. Zhang, P., Xu, T., Wang, S., Yang, X., Sun, P., Jia, P., Lin, J., Wang, B., Zhang, Y., Meng, D., et al. (2026). Highly accurate ab initio gene annotation with ANNEVO. Nature Methods 23:740–748. 10.1038/s41592-026-03036-7.

11. Zhang, W., Zhang, T., Wu, Y., and Jiang, J. (2012). Genome-Wide Identification of Regulatory DNA Elements and Protein-Binding Footprints Using Signatures of Open Chromatin in Arabidopsis. The Plant Cell 24:2719–2731. 10.1105/tpc.112.098061.

## Supplemental References

1. Gabriel, L., Becker, F., Hoff, K.J., and Stanke, M. (2024). Tiberius: end-to-end deep learning with an HMM for gene prediction. Bioinformatics 40:btae685.

2. Gabriel, L., Brůna, T., Kaur, A., Krishnan, A., Ortmann, F., Salamov, A., Talbot, S., Becker, F., Krieg, R., and Wheat, C.W. (2026). Accurate ab initio gene prediction in eukaryotes with Tiberius in multiple clades. bioRxiv.

3. Goodstein, D.M., Shu, S., Howson, R., Neupane, R., Hayes, R.D., Fazo, J., Mitros, T., Dirks, W., Hellsten, U., and Putnam, N. (2012). Phytozome: a comparative platform for green plant genomics. Nucleic acids research 40:D1178–D1186.

4. Holst, F., Bolger, A.M., Kindel, F., Günther, C., Maß, J., Triesch, S., Kiel, N., Saadat, N., Ebenhöh, O., Usadel, B., et al. (2026). Helixer: ab initio prediction of primary eukaryotic gene models combining deep learning and a hidden Markov model. Nature Methods 23:732–739. 10.1038/s41592-025-02939-1.

5. Hu, E.J., Shen, Y., Wallis, P., Allen-Zhu, Z., Li, Y., Wang, S., Wang, L., and Chen, W. (2021). Lora: Low-rank adaptation of large language models. arXiv preprint arXiv:2106.09685.

6. Lin, K., Zhang, Q., Wang, R., Hu, X., and Xu, W. (2025). PlantBiMoE: a bidirectional foundation model with SparseMoE for plant genomes. In 2025 IEEE International Conference on Bioinformatics and Biomedicine (BIBM). 10.1109/BIBM66473.2025.11356776.

7. Lin, T.-Y., Goyal, P., Girshick, R., He, K., and Dollár, P. (2017). Focal loss for dense object detection. Proceedings of the IEEE international conference on computer vision.

8. Linder, J., Srivastava, D., Yuan, H., Agarwal, V., and Kelley, D.R. (2025). Predicting RNA-seq coverage from DNA sequence as a unifying model of gene regulation. Nature Genetics 57:949–961. 10.1038/s41588-024-02053-6.

9. Loshchilov, I., and Hutter, F. (2017). Decoupled weight decay regularization. arXiv preprint arXiv:1711.05101.

10. Lukashin, A.V., and Borodovsky, M. (1998). GeneMark. hmm: new solutions for gene finding. Nucleic acids research 26:1107–1115.

11. O’Leary, N.A., Wright, M.W., Brister, J.R., Ciufo, S., Haddad, D., McVeigh, R., Rajput, B., Robbertse, B., Smith-White, B., Ako-Adjei, D., et al. (2016). Reference sequence (RefSeq) database at NCBI: current status, taxonomic expansion, and functional annotation. Nucleic Acids Res 44:D733–745. 10.1093/nar/gkv1189.

12. Pertea, G., and Pertea, M. (2020). GFF utilities: GffRead and GffCompare. F1000Research 9:ISCB Comm J-304.

13. Ronneberger, O., Fischer, P., and Brox, T. (2015). U-Net: Convolutional Networks for Biomedical Image Segmentation. In N. Navab and J. Hornegger and W.M. Wells and A.F. Frangi, eds. Medical Image Computing and Computer-Assisted Intervention – MICCAI 2015. Springer International Publishing.

14. Schiff, Y., Kao, C.H., Gokaslan, A., Dao, T., Gu, A., and Kuleshov, V. (2024). Caduceus: Bi-Directional Equivariant Long-Range DNA Sequence Modeling. Proc Mach Learn Res 235:43632–43648.

15. Shazeer, N., Mirhoseini, A., Maziarz, K., Davis, A., Le, Q.V., Hinton, G.E., and Dean, J. (2017). Outrageously Large Neural Networks: The Sparsely-Gated Mixture-of-Experts Layer. ArXiv abs/1701.06538.

16. Su, J., Ahmed, M., Lu, Y., Pan, S., Bo, W., and Liu, Y. (2024). Roformer: Enhanced transformer with rotary position embedding. Neurocomputing 568:127063.

17. Tegenfeldt, F., Kuznetsov, D., Manni, M., Berkeley, M., Zdobnov, E.M., and Kriventseva, E.V. (2025). OrthoDB and BUSCO update: annotation of orthologs with wider sampling of genomes. Nucleic Acids Research 53:D516–D522. 10.1093/nar/gkae987.

18. Touvron, H., Martin, L., Stone, K., Albert, P., Almahairi, A., Babaei, Y., Bashlykov, N., Batra, S., Bhargava, P., and Bhosale, S. (2023). Llama 2: Open foundation and fine-tuned chat models. arXiv preprint arXiv:2307.09288.

19. Vaswani, A., Shazeer, N., Parmar, N., Uszkoreit, J., Jones, L., Gomez, A.N., Kaiser, Ł., and Polosukhin, I. (2017). Attention is all you need. Advances in neural information processing systems 30.

20. Zhang, P., Xu, T., Wang, S., Yang, X., Sun, P., Jia, P., Lin, J., Wang, B., Zhang, Y., Meng, D., et al. (2026). Highly accurate ab initio gene annotation with ANNEVO. Nature Methods 23:740–748. 10.1038/s41592-026-03036-7.

